# LDL Signals Through Scarf1 to Drive a Pro-Fibrotic Secretory Response in Endothelial Cells

**DOI:** 10.1101/2025.08.15.670628

**Authors:** Xiangyi Gan, Salem Alamri, Harry Mellor

## Abstract

**BACKGROUND:** Endothelial cells lining the vasculature form a polarised secretory organ that releases proteins apically into the circulation and basally into the subendothelial matrix. While this secretion plays many critical roles in homeostasis, the potential roles for endothelial secretion in vascular disease remain under-studied.

**METHODS:** We treated polarised monolayers of primary human endothelial cells with native and oxidised LDL and identified the proteins secreted from the apical and basal endothelial surfaces by tandem mass tag mass spectrometry.

**RESULTS:** Treatment with either native or oxidised LDL led to increased secretion of a cohort of 21 mostly pro-fibrotic proteins. We focussed on fibronectin, which was secreted apically or basally, depending on the direction of LDL treatment. LDL-stimulated fibronectin secretion did not require well-characterised endothelial LDL receptors but was instead mediated by the poorly characterised scavenger receptor Scarf1 and the cholesterol-sensitive SNARE proteins syntaxin 4 and 6. This LDL-stimulated secretion did not involve increased fibronectin expression but instead appeared to result from a decrease in fibronectin turnover and a re-routing of intracellular sorting.

**CONCLUSIONS:** This novel endothelial secretory pathway links circulating LDL to the release of pro-fibrotic proteins from the endothelium, supporting a role for endothelial cell secretion in the progression of lipid-induced fibrotic disease.

## Introduction

The endothelium acts as a selective barrier between the circulation and the body, controlling the transport of fluid, macromolecules and cells between these two compartments. It is also a major secretory organ, releasing factors to regulate vascular tone, thrombosis, haemostasis and inflammation. ^1,2^ In an adult human, the endothelium comprises 10-60 trillion endothelial cells (ECs) and weighs approximately 1kg, rivalling the liver in mass and making it one of the largest secretory organs in the body. ^3^ Endothelial secretion is tissue-specific, with ECs secreting different combinations of angiocrine factors to mediate local regulation of tissue homeostasis and regeneration. ^4,5^

EC secretion is polarised, allowing secretion of chemokines and clotting factors directly into the circulation from the apical/luminal surface ^6,7^. For example, the proinflammatory chemokines IL-6 and TNF-α are routed through intracellular trafficking pathways to the apical surface of ECs, allowing these mediators to reach more distant tissues via the blood. ^8^ In addition to the secretion of soluble proteins, ECs release membrane bound extracellular vesicles to carry long-range signals to distant tissues via the circulation. ^9^ ECM proteins and regulators are secreted from the basal/abluminal surface, allowing ECs to actively modulate their local tissue environment ^5^. Examples include perlecan, a major component of the endothelial basement lamina, and thrombospondin-1, a matricellular protein that acts as a brake on angiogenic signalling. ^10^ Additional factors are secreted basally to communicate with perivascular cells. Basal secretion of PDGFβ regulates mural cell recruitment and proliferation. ^11^ The secretion of endothelin-1 is highly polarized to the basal surface, acting on vascular smooth muscle cells in the vessel wall to drive vasoconstriction. ^12^ Interestingly, some proteins can be released from either surface, depending on context. For example, ECs constitutively secrete von Willebrand Factor as a low molecular weight form from their basal surface at rest, but as a multimerized protein from their apical surface upon activation. Basally-secreted von Willebrand Factor provides a reservoir of collagen-bound subendothelial protein, whereas apically-secreted multimerized factor promotes platelet binding and thrombosis ^13^.

Previously, we constructed a proteomic platform to allow study of polarised EC secretion in depth. ^10^ ECs are cultured as quiescent monolayers on Transwell cell culture inserts, allowing collection of media from the apical and basal faces separately (Figure 1A). Cells are grown in a specially formulated media; HIFA2, which is protein-free and optimised for barrier function. ^10^ This platform allowed us to characterise the polarised EC secretome for the first time, showing that it is possible to obtain deep (c.a. 1,000 proteins) secretomes from relatively small (c.a. 100,000) numbers of primary human ECs. ^10^ Given the importance of polarised endothelial secretion to normal functioning of the endothelium, we were interested in investigating if dysregulation of these pathways contributes to vascular disease.

**Figure 1.**
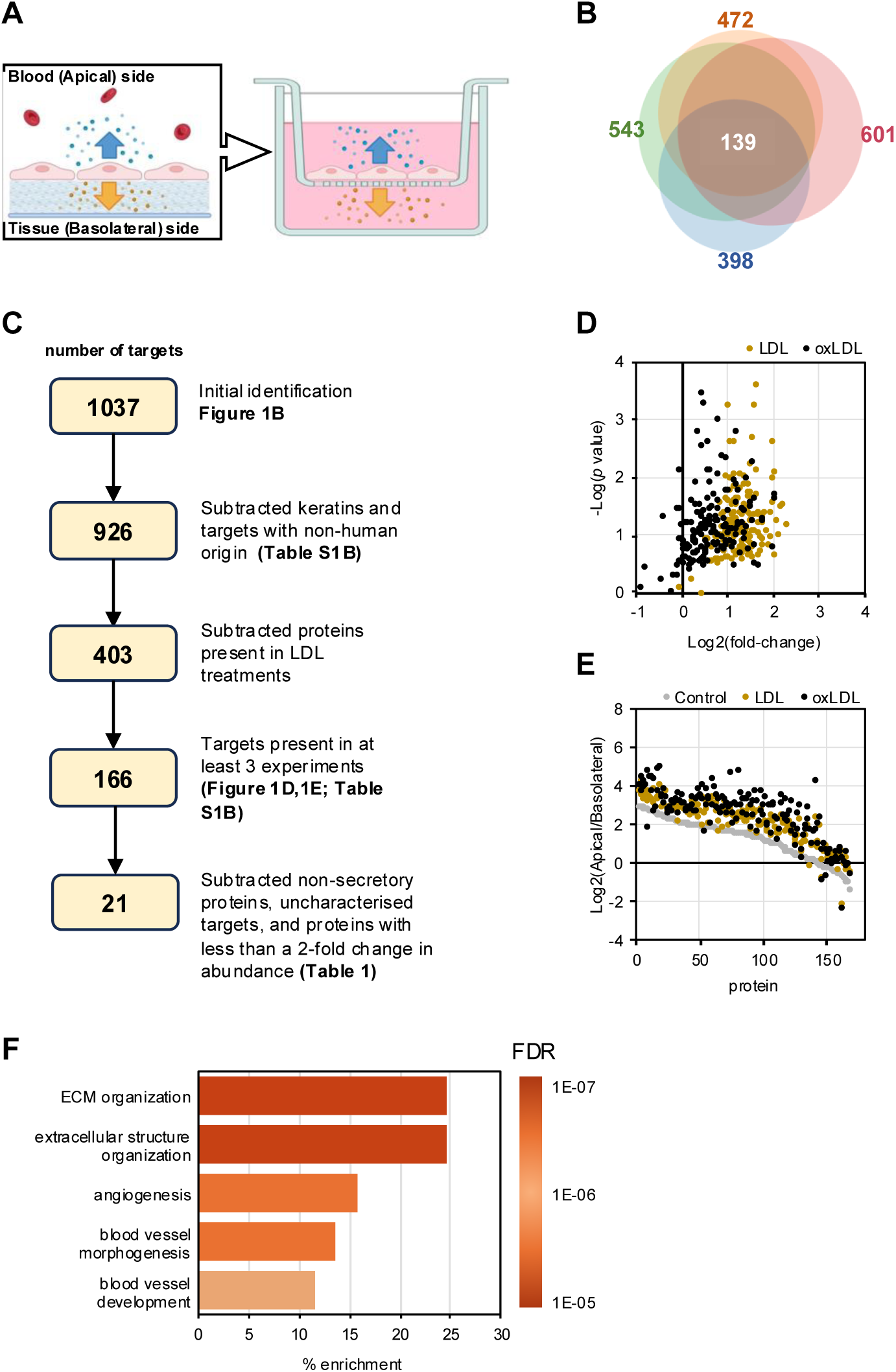
Endothelial Cells Produce a Secretory Response to Oxidised and Native LDL. **A**, Illustration of the Transwell system. **B**, ECs were grown to confluence in Transwell inserts and then transferred to HIFA2 medium. The cells were treated from the apical side with 50µg/mL of nLDL or oxLDL for 48h and the conditioned media were collected and analysed by TMT-mass spectrometry. Data represent the averages of four independent experiments. The Venn diagram shows the total number of proteins detected in each experimental repeat and the overlaps in identification between the datasets. **C,** A flow chart of the filtering steps used and the number of targets remaining at each step. **D**, The volcano plot shows a comparison of nLDL and oxLDL-induced total secretion changes (apical + basal abundance). Log2 of the fold-changes in abundance were plotted against -Log10 of the corresponding *P* values for the 166 proteins. **E**, The plot shows changes in the polarity of secretion of the 166 proteins upon apical treatment with nLDL or oxLDL, compared to control. Log2 of the secretion polarity ratio (apical/basolateral abundance) was compared to the control samples (grey) for each protein. **F**, Gene ontology analysis of the final candidate list of 21 secreted proteins shows enrichment for ECM components and regulators of angiogenesis. The colour scale shows the false discovery rate (FDR).

Previous work has shown that oxidised low-density lipoprotein (oxLDL) alters the secretion of fibronectin by ECs. ^14^ Fibronectin is an ECM protein normally secreted from the basal surface of ECs ^15,16^. On treatment with minimally-modified LDL (mmLDL) or oxLDL, ECs accumulate patches of fibronectin on their apical surface. These unnatural adhesive patches promote monocyte binding to the EC surface. ^14^ Quiescent ECs normally exhibit low levels of fibronectin secretion, but this is upregulated during angiogenesis, contributing to a provisional matrix that supports neovessel growth. ^17^ ECs secrete alternatively spliced forms of cellular fibronectin (cFn), which contain EIIIA/EDA and/or EIIIB/EDB domains. ^18,19^ This fibronectin is normally deposited into the subendothelial matrix and plays a critical role in vessel formation during development. ^18^ Fibronectin expression commences during vasculogenesis and continues throughout angiogenesis. ^20^ In retinal angiogenesis, endothelial tip cells follow tracks of fibronectin laid down by astrocytes ^21^; however, ECs also secrete fibronectin themselves, which acts as a cell autonomous autocrine signal. ^22^ Fibronectin binds VEGF in the provisional matrix and presents it to tip cells ^21^. It also interacts directly with integrins on the endothelial cell surface to drive alignment of the cytoskeleton and to promote 3D vessel formation ^23^. Cell-specific targeting of fibronectin expression shows that the fibronectin secreted by ECs has non-redundant roles in vessel sprouting and vessel stability. ^22^ Endothelial fibronectin expression decreases at the end of angiogenesis, when the provisional matrix is replaced with the basement lamina and the vessels become stable ^24^.

Interestingly, endothelial fibronectin expression is also upregulated in the early stages of atherosclerosis. Fibronectin accumulation is seen in fatty streaks and in early atherosclerotic lesions, but is absent from the surrounding unaffected tissue. ^25^ This fibronectin expression is triggered by atherogenic flow patterns and contributes to local inflammatory signalling. ^26,27^ Genetic deletion of the EIIIA/EDA fibronectin isoform in mouse models of atherosclerosis leads to reduced lesion size, supporting a direct role for cFn in disease progression. ^28^ Genetic models show that while fibronectin in mature plaques is secreted by vascular smooth muscle cells, in early plaques fibronectin is secreted by ECs. In the absence of EC fibronectin, initiation of atherosclerosis is inhibited with reduced infiltration of inflammatory cells. ^29^ Fibronectin expression is also upregulated in fibrosis, where it mediates the deposition of collagen I and other pro-fibrotic ECM proteins. ^30^

Fibronectin plays an essential role in the development of pulmonary and hepatic fibrosis. ^31,32^ While the primary source of fibronectin in fibrosis is myofibroblast cells, fibronectin expression is also upregulated in liver sinusoidal ECs in early hepatic fibrosis and in lung ECs in pulmonary fibrosis. ^32,33^ Increases in fibronectin in the subendothelial space in fibrosis can lead to disruption of barrier function and conversion of ECs into myofibroblasts – further contributing to fibrosis. ^34^

We were interested in further investigating the polarised secretion of fibronectin by ECs in response to oxidised LDL. We hypothesised that this may be part of a broader response, and we set out to characterise that secretome and to investigate the regulation of its secretion.

## METHODS

### Data Availability

All supporting data for this study are available from the corresponding author on request. For detailed experimental methods, materials, and statistical analyses, please see the Supplemental Material and Major Resource Table.

The following citations are for the expanded Materials and Methods and Major Resource Table sections. ^10,73–75^

## RESULTS

### Endothelial Cells Produce a Secretory Response to Oxidised and Native LDL

ECs were cultured to confluence on semi-permeable Transwell filters (Figure 1A) to allow the formation of a polarised endothelial monolayer, before switching to HIFA2 medium. Preliminary assessments showed LDL treatments did not disrupt the barrier function of the endothelial layer, ensuring minimal protein leakage to the basal compartment (Figure S1C-D). ECs were treated with 50 µg/mL of native LDL (nLDL) or oxLDL for 48h, and conditioned media from the apical and basal chambers of the Transwell system were then collected separately for analysis by tandem mass tag (TMT) mass spectrometry. We also analysed the nLDL and oxLDL preparations alone to characterise their protein content and account for it in the treated samples (Table S1A, Figure S1E). Secretome data were collected from 4 independent experiments. After removal of contaminants and the serum proteins found in the LDL preparations, we identified 166 proteins present in at least 3 experiments (Table S1B, Figure 1C). Treatment with either nLDL or oxLDL led to significantly increased secretion of a subset of EC proteins (Figure 1D). Unexpectedly, their secretion was triggered by both LDL species (Figure 1D). We analysed the polarity of secretion and found that lipoprotein treatment affected this strongly, with a general rerouting from basolateral to apical secretion (Fig 1E). We refined the target list by filtering for proteins whose secretion increased by at least 2-fold in response to LDL. We also removed proteins that are not known to be secreted (Table S1B), leaving us with a candidate list of 21 secreted proteins. Gene ontology analysis showed that the list was enriched for ECM proteins and regulators of angiogenesis and blood vessel development (Figure 1F). The list of 21 high-confidence hits is shown in Table 1. In general, nLDL had a slightly greater effect on secretion than oxLDL. Surprisingly, in all cases, treatment with lipoprotein promoted rerouting of basally secreted ECM proteins to the apical surface. To summarise, treatment with either nLDL or oxLDL led to increased secretion of a cohort of 21 proteins and switched the polarity of their secretion towards apical release.

**Table 1.** Proteins secreted in response to LDL. The table shows the 21 proteins identified as secreted in response to apical treatment of ECs with nLDL and oxLDL. The fold-change (FC) in secretion is given, as is the polarity ratio of apical:basal secretion. In general, proteins showed increased secretion on treatment with nLDL, compared to oxLDL. Treatment with LDL from the apical surface generally caused a switch in polarity of secretion towards more apical release. Data are means of n=3 independent experiments.

| Gene | Protein | FC over control |  | Apical/basal ratio |  |  |
| --- | --- | --- | --- | --- | --- | --- |
|  |  | LDL | oxLDL | Control | LDL | oxLDL |
| SERPINE1 | PAI-1 | 3.70 | 2.73 | 1.61 | 3.52 | 4.91 |
| <u>MMP1</u> | MMP1 | 3.54 | 3.32 | 1.65 | 4.16 | 6.23 |
| CCN2 | CCN2 | 3.20 | 2.00 | 2.50 | 2.10 | 1.10 |
| NAMPT | NAMPT | 3.20 | 1.13 | 3.90 | 13.28 | 7.53 |
| PTX3 | Pentraxin-related protein 3 | 3.09 | 1.61 | 1.52 | 2.55 | 3.02 |
| CLU | Clusterin | 3.07 | 1.90 | 0.74 | 1.50 | 1.85 |
| <u>CCN1</u> | CCN family member 1 | 3.06 | 2.15 | 1.18 | 2.15 | 3.44 |
| LAMC1 | Laminin subunit gamma-1 | 2.83 | 0.90 | 0.81 | 1.55 | 1.17 |
| PXDN | Peroxidasin | 2.81 | 1.60 | 2.29 | 3.69 | 4.26 |
| FSTL1 | Follistatin-related protein 1 | 2.77 | 1.32 | 0.80 | 1.27 | 1.63 |
| GRN | Progranulin | 2.69 | 2.19 | 1.21 | 2.02 | 2.17 |
| QSOX1 | Sulfhydryl oxidase 1 | 2.58 | 0.97 | 1.45 | 2.12 | 2.00 |
| EFEMP1 | EFEMP1 | 2.46 | 1.31 | 0.90 | 1.31 | 1.10 |
| LTBP2 | LTBP-2 | 2.43 | 2.19 | 1.59 | 3.09 | 3.31 |
| IGFBP7 | IGF binding protein 7 | 2.43 | 1.67 | 0.88 | 1.48 | 1.56 |
| <u>FN1</u> | Fibronectin 1 | 2.39 | 1.77 | 0.71 | 1.51 | 1.59 |
| TIMP2 | Metalloproteinase inhibitor 2 | 2.34 | 1.19 | 0.70 | 1.64 | 1.26 |
| NPC2 | NPC2 | 2.18 | 1.00 | 1.16 | 1.42 | 1.54 |
| LOXL2 | Lysyl oxidase homolog 2 | 2.18 | 1.48 | 0.83 | 1.42 | 1.26 |
| THBS1 | Thrombospondin-1 | 2.15 | 1.81 | 0.61 | 1.33 | 1.59 |
| TCN2 | Transcobalamin-2 | 2.10 | 1.38 | 0.92 | 1.62 | 2.12 |

### Endothelial Cells Secrete a Cohort of Pro-Fibrotic Proteins in Response to LDL

Closer inspection allowed for a clearer picture of these secreted proteins (Table 1). The secretome is enriched in proteins that drive pro-fibrotic remodelling and early atherogenic change. CCN2 is a highly profibrogenic matricellular protein that is overexpressed in fibrotic lesions. ^35^ In fibrotic liver tissue CCN2 is produced by activated hepatic stellate cells, fibroblasts and endothelial cells, promoting collagen synthesis and stellate cell activation.

PAI-1 is upregulated in many chronic liver diseases and drives fibrosis by inhibiting matrix degradation. ^36^ Genetic loss of PAI-1 in mice significantly mitigates liver fibrosis ^37^ whereas PAI-1 overexpression correlates with exacerbated scar formation. Lysyl oxidase 2 covalently crosslinks collagen fibres, stabilizing fibrotic scars. ^38^ Timp-2 inhibits collagen degrading enzymes in the ECM to stabilise collagen deposition. ^39^ Its levels rise in activated hepatic stellate cells and fibrotic liver, tipping the balance toward collagen accumulation.

Thrombospondin-1 activates latent TGF-β, one of the primary drivers of fibrotic responses. ^40^ LTBP2 is upregulated in fibrosis, driving activation of hepatic stellate cells in liver fibrosis ^41^ and myocardial fibrosis in dilated cardiomyopathy. ^42^ Atherosclerosis shares several regulators with fibrosis and can be viewed as a localised fibrotic disease. PAI-1 ^43^, MMP1 ^44^, pentraxin-3 ^45^ and thrombospondin-1 ^46^ have all been shown to contribute to the progression of atherosclerosis, with overlapping and/or related roles to those in fibrosis. Taken together, the proteins released by ECs in response to LDL constitute a pro-fibrotic cocktail, focussed on matrix remodelling and the amplification of TGF-β-driven pathways.

### LDL Directs Polarised Secretion in Endothelial Cells

From the list of identified proteins, fibronectin, CCN1, and MMP1 were selected for further investigation based on their functional roles and potential involvement in lipoprotein-induced endothelial dysfunction. To validate the proteomic findings and to quantify the secretion of the three proteins, ELISA assays were performed using conditioned media collected from the Transwell system, following the same experimental protocol as the proteomic analysis. As previously, we treated endothelial monolayers with lipoprotein from the apical side to mimic exposure to circulating LDL. Prolonged exposure to a high-fat diet can lead to accumulation of LDL in the subendothelial space. This LDL becomes oxidised leading to the generation of pro-atherogenic signals that are seen by the basal surface of the endothelial cells. ^47^ To examine the effects of basally-presented LDL, we also treated monolayers from the basal side. Apical treatment with nLDL led to a significant increase in the apical secretion of fibronectin, MMP1 and CCN1, confirming the proteomic data (Figure 2A-C). Intriguingly, basal treatment with nLDL led to a greater increase in fibronectin secretion and this was now secreted basally (Figure 2A). A similar result was obtained with MMP1 secretion (Figure 2B). With CCN1 secretion, basal nLDL treatment stimulated apical release, although at a much lower level than achieved by apical treatment (Figure 2C). Essentially identical results were obtained with oxLDL (Figure 3D-F). These findings show that for the secretion of fibronectin and MMP1, the direction of the LDL signal determines the direction of secretion.

**Figure 2.**
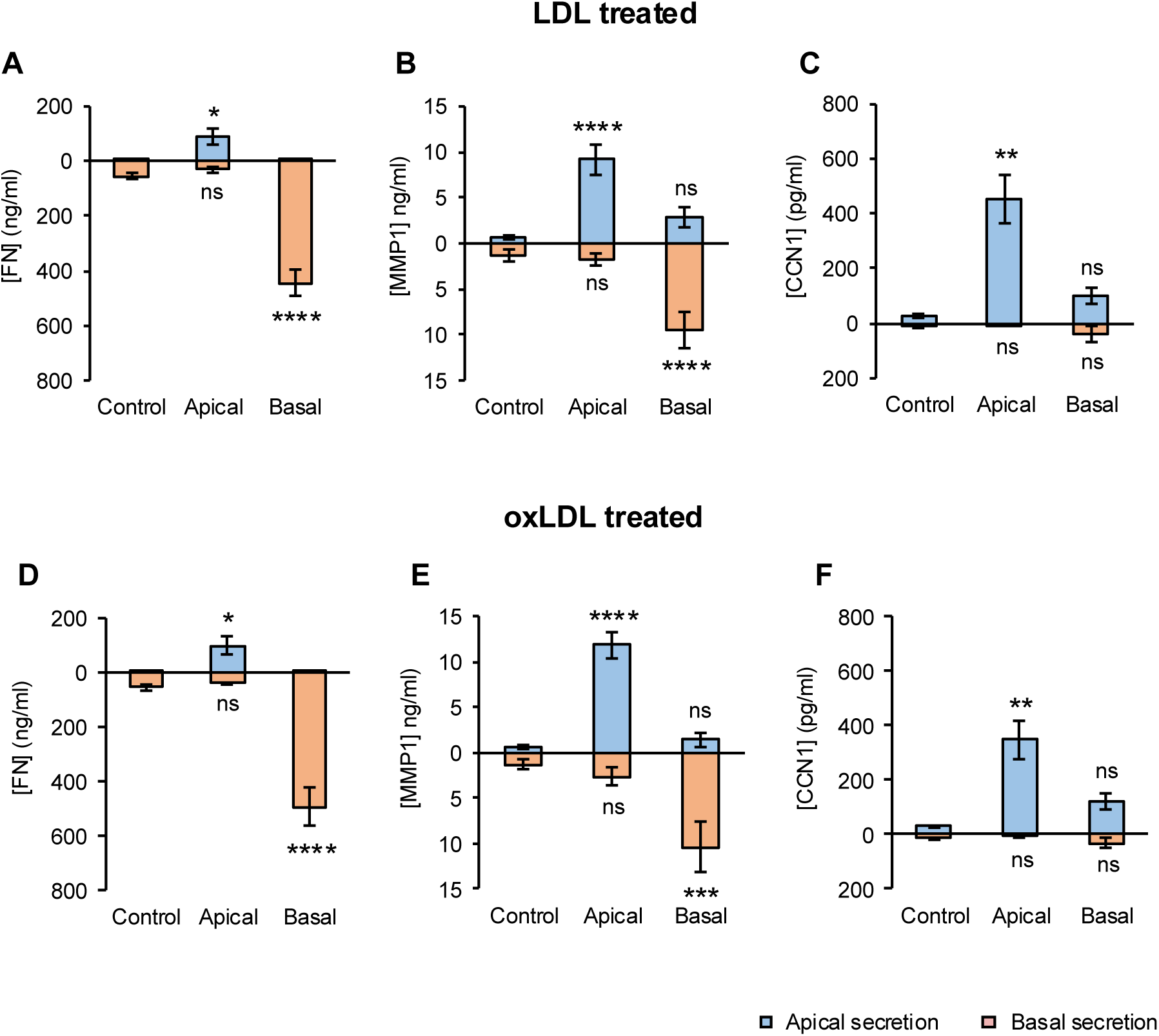
LDL Directs Polarised Secretion in Endothelial Cells. ECs were cultured to confluence on Transwell inserts. 50µg/mL of nLDL (**A-C**) or oxLDL (**D-F**) was administrated from either the apical or the basal side. The direction of treatment is indicated on the x-axis. After 48h, the conditioned media were collected, and the concentrations of apically secreted proteins (blue bars) or basally secreted proteins (orange bars) were measured for each target by ELISA. For fibronectin and MMP1, the direction of treatment drove the direction of secretion. Data are means ± SEM; n = 3 experiments.

**Figure 3.**
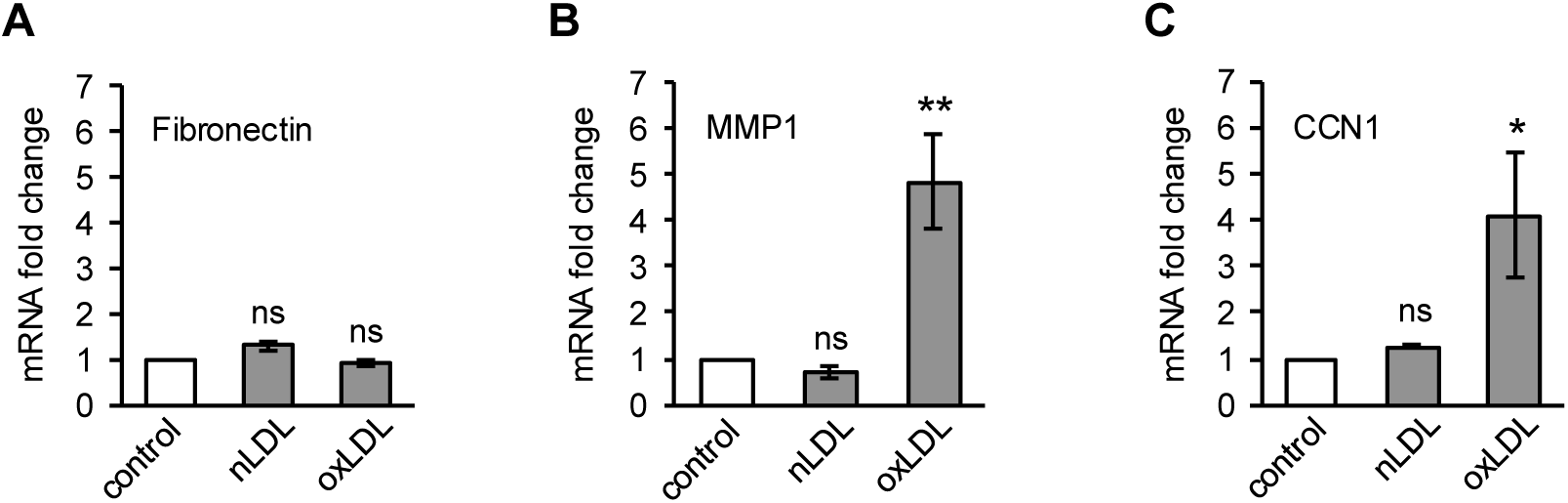
LDL-induced fibronectin secretion is regulated post-transcriptionally. ECs were grown to confluence and treated with 50µg/mL of LDL or oxLDL for 48h as before. The mRNA levels of **A,** fibronectin, **B,** MMP1, and **C,** CCN1 were measured by Q-PCR. Data are means ± SEM; n = 3 experiments.

### LDL Drives Fibronectin Secretion by Stabilising Fibronectin Turnover

Previous work has shown that oxLDL can increase the gene expression of MMP1 in ECs and CCN1 in macrophages, both at the transcriptional level ^48,49^. To investigate whether the increased secretion of fibronectin, MMP1 and CCN1 was driven by increased expression, we measured mRNA levels of each target. As reported previously, oxLDL stimulated the expression of MMP1 and CCN1 (Figure 3B, C); however, fibronectin expression was not affected. Native LDL had no effect on the expression of any target (Figure 3A-C). We conclude that transcriptional regulation may contribute to the effects of oxLDL on secretion of MMP1 and CCN1; however, there must be additional mechanisms to explain the effects of nLDL on endothelial secretion, and to explain increased fibronectin secretion triggered by either native or oxidised LDL.

To investigate this, we focussed on the turnover of fibronectin. ECs secrete cFn basally, where it is assembled into fibronectin fibrils in the vascular matrix. ^15,16^ ECs remodel these fibrils by re-internalising the fibronectin, trafficking it through endosomal recycling pathways and re-depositing it as new ECM. ^50,51^ One way for ECs to rapidly secrete fibronectin without new protein synthesis would be to release fibronectin stored in the ECM by endocytosing and re-secreting this protein. To explore this, we treated ECs with native or oxidised LDL and measured the density of fibronectin fibres in the ECM by immunofluorescence microscopy. We used an antibody specific for EDA-fibronectin; the major cellular isoform secreted by endothelial cells. ^52^ Figure 3E shows that treatment with either native or oxidised LDL led to a significant increase in fibronectin fibre density. We carried out deoxycholate (DOC) solubility assays to confirm this. In this assay, cells are extracted in DOC and the insoluble ECM fibrils separated from the soluble cellular cFn pool, before analysis by western blotting. ^53^ Treatment with either nLDL or oxLDL caused a significant increase in both deposited fibronectin fibrils and the cellular pool of soluble fibronectin (Figure 3F). We conclude that pre-deposited fibrillar fibronectin is not the net source of secreted fibronectin in LDL treated ECs.

The cyclic processes of fibronectin reinternalization and recycling mean that there is a significant flux of fibronectin through the endosomal compartment. Although most internalised fibronectin is recycled, a fraction is degraded in the lysosome. ^54,55^ We hypothesised that LDL treatment might trigger rerouting of fibronectin intracellular traffic to spare it from lysosomal degradation, so increasing the pool of protein available for secretion. This would explain the increase in the cellular pool seen in Figure 3E. To examine constitutive fibronectin turnover in ECs, we applied cycloheximide to inhibit new fibronectin synthesis. Treatment with cycloheximide for 24h led to an almost complete loss of both secreted fibronectin and deposited fibrillar fibronectin (Figure 4A), demonstrating the constitutive degradation of the ECM pool. We treated ECs with Bafilomycin A1 to block trafficking of endocytosed proteins to the lysosome. This led to a loss of deposited fibronectin fibrils but had no effect on fibronectin secretion (Figure 4A). Confocal slices of the body of the cells showed that the internalised fibronectin had accumulated in cytosolic vesicles, in keeping with a block in onward trafficking of endocytosed fibronectin. Finally, we incubated ECs with pepstatin, which inhibits lysosomal proteases without blocking endosomal trafficking. This led to a significant increase in constitutive fibronectin secretion, with maintenance of deposited fibrillar fibronectin (Figure 4D).

**Figure 4.**
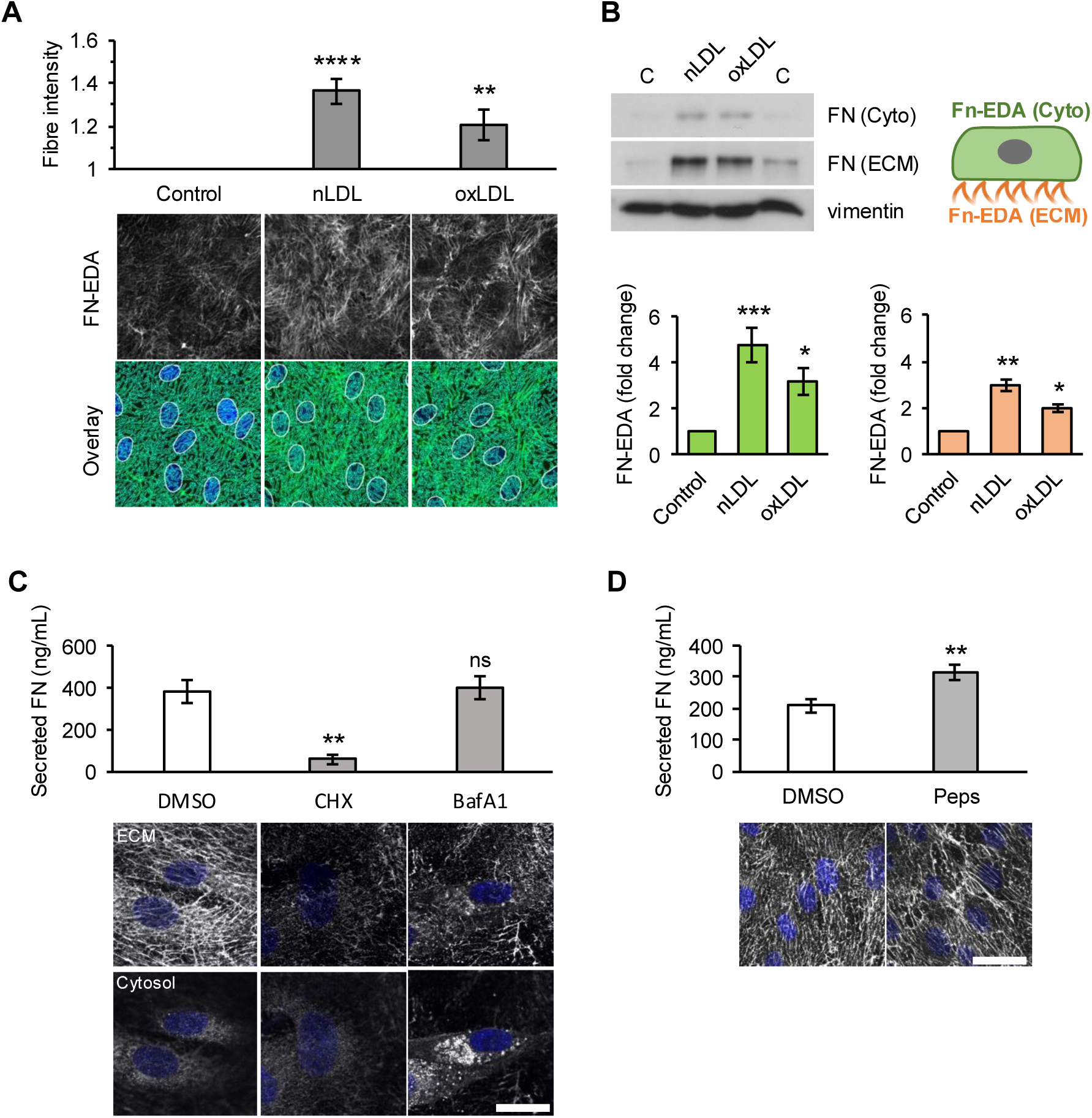
LDL-induced fibronectin secretion is supported by reduction of intercellular turnover. **A,** ECs were grown to confluence on coverslips and treated with 50µg/mL of nLDL or oxLDL for 48h. Cells were fixed and stained for EDA fibronectin and fibronectin fibre intensity was quantified in ImageJ. Data are means ± SEM; n = 4 experiments. The top panels show representative cell images from the analysis. The bottom panels show the extracted features. Scale bar = 20μm. **B,** ECs were treated with nLDL or oxLDL as before and then extracted in 1% sodium deoxycholate (DOC) to separate the insoluble matrix from the cytosolic pool. The lysates were then analysed by western blotting for EDA fibronectin. The DOC assays were quantified by densitometry. Treatment with either nLDL or oxLDL increased fibronectin deposition in both assays (**A,B**). **C,** ECs were treated with DMSO, 35μM cycloheximide (CHX), or 50nM bafilomycin A1 (BafA1) for 24h. Secreted fibronectin was measured in the cell media by ELISA, and the cells were fixed and stained for EDA fibronectin. Confocal slices are shown at the level of the ECM and the cell cytosol. Inhibition of new fibronectin synthesis with CHX led to a loss of both deposited fibronectin fibrils and secreted soluble fibronectin. Treatment with BafA1 led to a loss of deposited fibronectin and an accumulation of fibronectin in cytosolic vesicles. **D,** ECs were treated with DMSO or 2µg/mL pepstatin for 24h. Secreted fibronectin was measured in the cell media by ELISA, and the cells were fixed and stained for EDA fibronectin. Inhibition of lysosomal proteolysis with pepstatin led to a significant increase in fibronectin secretion, and a small increase in deposition. Data are means ± SEM; n = 3 experiments. Scale bars = 10 μm.

We conclude that constitutive degradation of fibronectin acts as a post-translational control mechanism to limit its secretion from ECs. We propose that LDL treatment reduces this constitutive turnover in ECs, leading to increased fibronectin secretion. When this fibronectin is released basally, integrins are able to assemble it into deposited fibrils, ^27^ whereas apical release results in predominately soluble secretion into the circulation. While we did not observe the apical patches of fibronectin seen by Shih *et al.* in oxLDL treated ECs, ^14^ those may have resulted from capture of apically-released fibronectin by apically localised integrins in their system. ^56^

### SCARF1 Mediates LDL-Induced Fibronectin Secretion

LDL receptor (LDLR) is the major cellular receptor for nLDL. ^57^ LDLR binds nLDL particles at the cell surface and is internalised via clathrin-dependent endocytosis. Acidification of the early endosomes leads to dissociation of nLDL, which is then trafficking to the lysosome, and LDLR is recycled to the cell surface. LDLR has a low affinity for oxLDL ^57^; however, endothelial cells also express scavenger receptors, which show specificity for oxLDL and trigger pro-inflammatory signals. In order to determine which LDL receptor(s) were responsible for the LDL-mediated secretory response, we silenced LDLR and the 3 best-characterised endothelial oxLDL receptors; LOX-1, SR-B1 and CD36. ^58^ The effectiveness of silencing was confirmed by Q-PCR (Figure S2A). Surprisingly, none of these receptors were required for the stimulation of fibronectin secretion by either nLDL or oxLDL. Silencing of LOX-1 in fact led to a small but significant increase in the response to both nLDL and oxLDL (Figure 5A).

**Figure 5.**
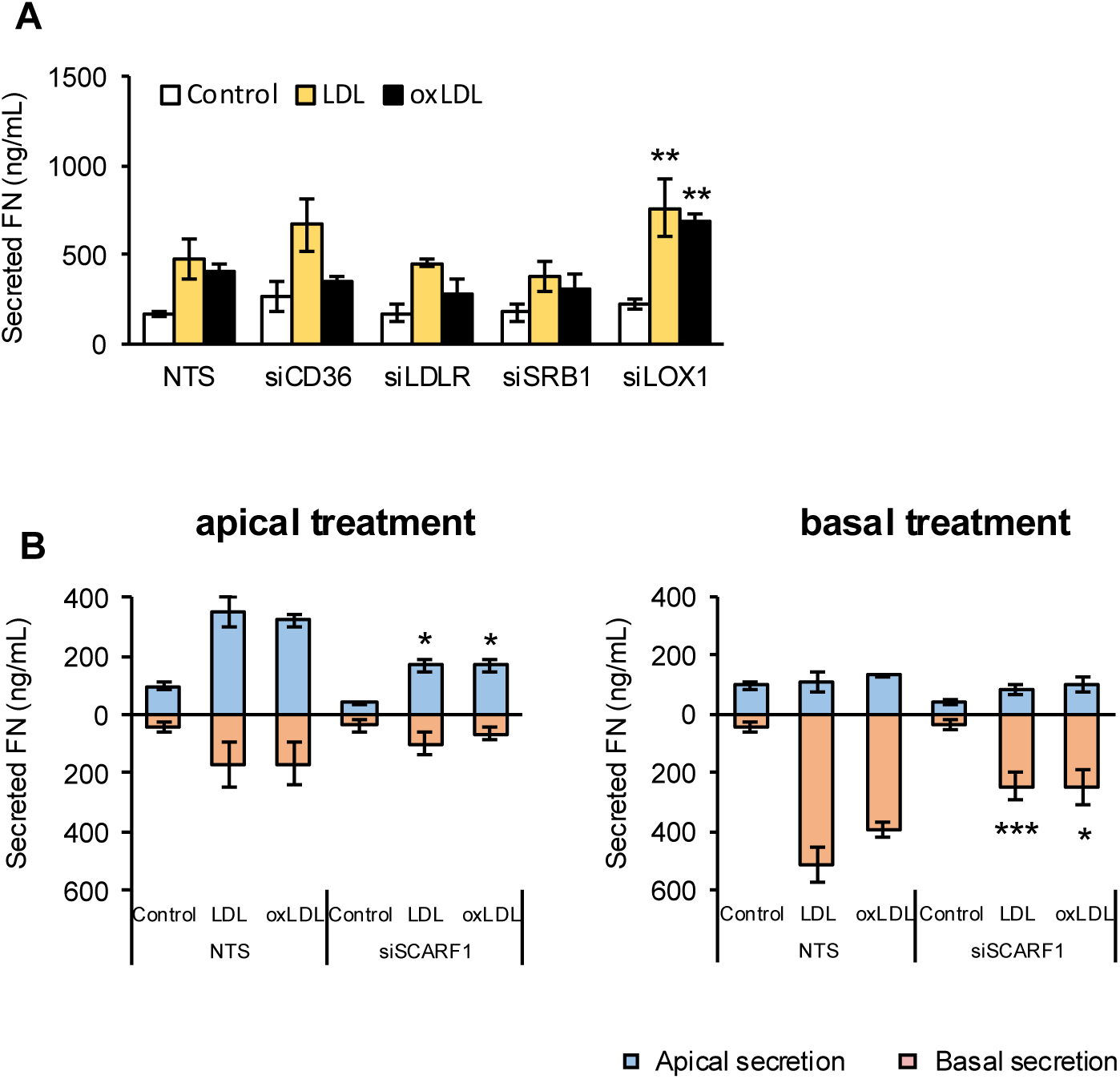
Scarf1 regulates LDL-induced fibronectin secretion. **A,** ECs were transfected with siRNAs targeting CD36, LDLR, SRB1, LOX1 or a non-targeting siRNA control (NTS) and grown to confluence on tissue culture dishes. The cells were treated with 50µg/mL of nLDL or oxLDL for 48h and the secreted fibronectin was measured by ELISA. Silencing of LOX1 led to a small but significant increase in fibronectin secretion in response to nLDL or oxLDL. **B,C,** ECs were transfected with siRNAs targeting Scarf1 or a non-targeting siRNA control and grown to confluence on Transwell inserts. 50µg/mL of nLDL or oxLDL was administrated from either the apical side of the Transwell (**B**) or the basal side (**C**). After 48h, the conditioned media were collected, and the concentrations of apically secreted (blue bars) or basally secreted fibronectin (orange bars) were measured by ELISA. Silencing of Scarf1 significantly reduced fibronectin secretion in response to nLDL or oxLDL from either treatment direction. Data are means ± SEM; n = 3 experiments.

In order to expand our search for the relevant receptor, we turned to a screening approach. ECs were treated with nLDL or oxLDL for either 30min or 24h. We used surface biotinylation to label all cell surface proteins and captured them on streptavidin-agarose. The samples were analysed by TMT mass spectrometry to identify proteins whose surface expression changed on LDL treatment (Table S2). Many cell surface receptors show altered surface expression in response to ligand binding. This may be short term change caused by internalisation through endocytosis, or longer-term desensitisations caused by degradation of the internalised receptor in the lysosome. LDLR is known to constantly recycle between the surface and the endosomal compartment and is not targeted for lysosomal degradation ^57^; however, treatment with nLDL over 24h leads to downregulation of receptor expression at the mRNA level. ^59^ In keeping with this, we saw no change in surface LDLR at 30min of treatment with nLDL, but a 57% decrease in surface expression at 24h (Table S2). Of the 541 surface proteins identified in the screen, only one other LDL receptor showed altered surface expression on LDL treatment: Scarf1 (SREC1; scavenger receptor expressed by endothelial cells 1). The surface level of Scarf1 increased in response to oxLDL treatment (98% at 30min, 28% at 24h; Table S2). This simple screen provided no definitive data on the trafficking of these receptors but suggested that Scarf1 was worth further investigation.

Scarf1 is a poorly-characterised scavenger receptor. ^60^ It was first identified in ECs, where it was shown to be a receptor for acetylated LDL ^61^. It has subsequently been shown to bind multiple diverse ligands, with roles in innate immunity and apoptotic cell clearance. ^60^ We silenced Scarf1 expression (Figure SB) and measured polarised fibronectin secretion in response to lipoprotein treatment. Silencing of Scarf1 significantly inhibited the fibronectin secretion in response to both nLDL and oxLDL, and this was the case for both apical and basal treatment (Figure 5B). We conclude that the endothelial secretory response to LDL is driven by Scarf1, and that it is separate to both the normal uptake of LDL by LDLR, and to previously studied endothelial scavenger receptor responses to oxLDL.

### LDL-Induced Fibronectin Secretion Involves the Cholesterol-Sensing Syntaxins, Stx4 and Stx6

We were interested in how binding of LDL to Scarf1 could drive changes in endothelial protein secretion and what signal mediated those changes. Scarf1 is not known to directly activate cell signalling pathways ^60^ and so we considered whether LDL itself may provide the signal. Uptake of nLDL by LDLR leads to trafficking of nLDL to the late endosome/lysosome, where cholesterol is released and trafficked onwards to multiple cell compartments, triggering several cellular signalling pathways. ^57^ This LDLR has a low affinity for oxLDL, meaning that oxLDL treatment does not lead to significant cholesterol uptake via LDLR. ^57^ Similarly, most scavenger receptors have a low affinity for nLDL and only increase cellular cholesterol when exposed to oxLDL. ^62^ We first tested whether treatment of ECs with native and oxidised LDL led to comparable levels of cholesterol loading – i.e., whether there was a common signal. We used Filipin staining of fixed cells to measure unesterified cellular cholesterol. ^63^ Incubation with either nLDL (Figure 6A) or oxLDL (Figure 6C) led to significant cholesterol uptake that was visible both at the plasma membrane and in intracellular vesicles. These vesicles co-localised strongly with the late endosomal marker CD63 (Figure 6A,B). Silencing of Scarf1 receptor expression significantly inhibited uptake of cholesterol in response to either nLDL or oxLDL (Figure 6C). We conclude that Scarf1 can mediate uptake of cholesterol from both native and oxidised LDL, and that this leads to trafficking of cholesterol through the endosomal pathway to the late endosome.

**Figure 6.**
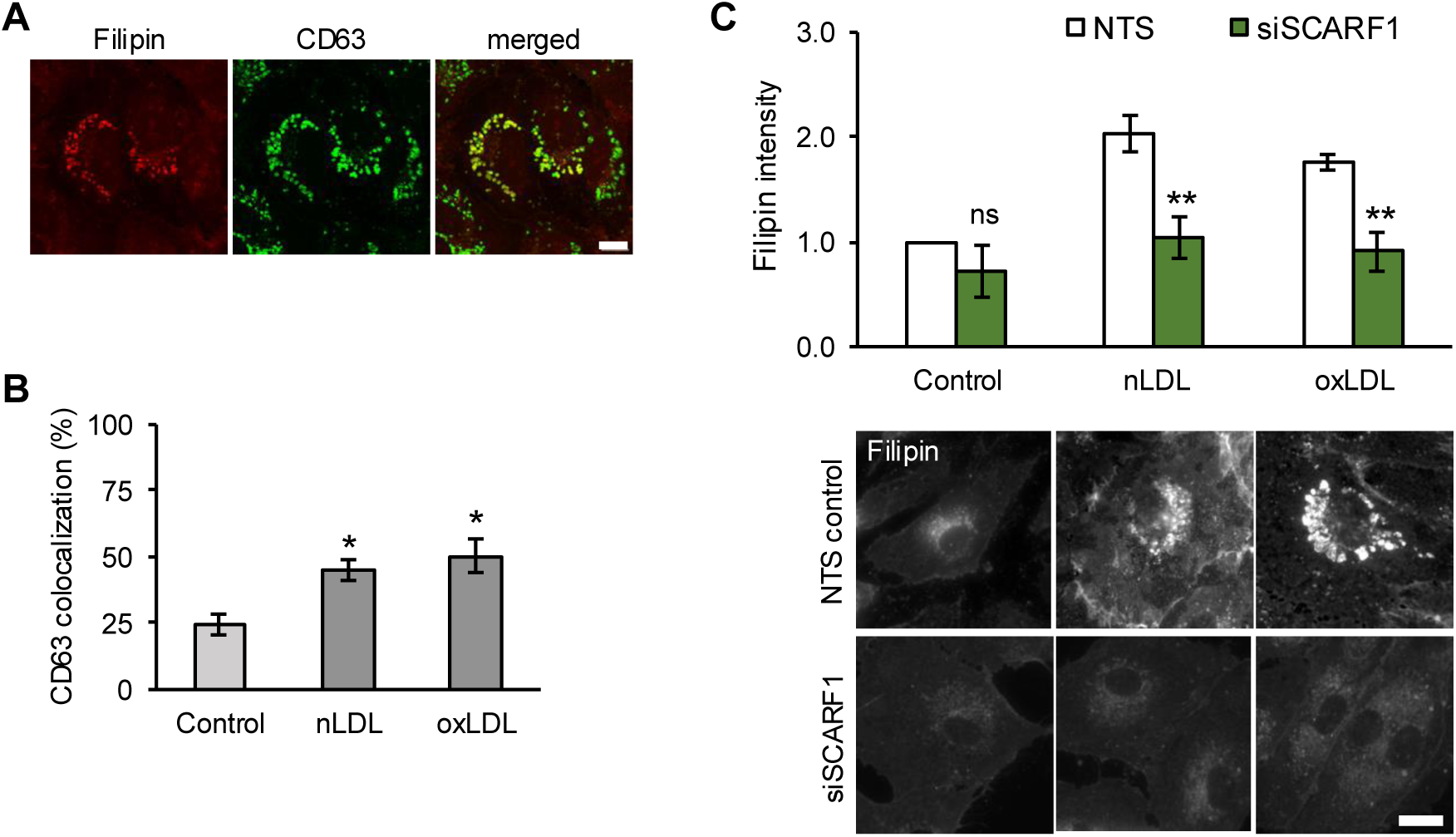
Scarf1 mediates the uptake of nLDL and oxLDL. **A.** ECs were treated with 50µg/mL of nLDL for 48h and then fixed and stained for CD63 (green) and with Filipin III to detect unesterified cholesterol (red). **B.** The percentage of colocalization between cholesterol+ vesicles and CD63 was determined from images of ECs treated for 48h with either 50µg/mL of nLDL or oxLDL. **C.** ECs were transfected with siRNAs targeting Scarf1or a non-targeting siRNA control (NTS) and grown to confluence on tissue culture dishes. The cells were treated with 50µg/mL of nLDL or oxLDL for 48h and then fixed and stained with Fillipin III. Uptake of cholesterol was quantified by measuring the intensity of Fillipin III staining. Silencing of Scarf1 significantly reduced cholesterol uptake from nLDL and oxLDL. Data are means ± SEM; n = 3 experiments. Scale bar = 10μm

SNARE proteins are key regulators of membrane trafficking and protein secretion. ^64^ Several SNAREs have been shown to bind cholesterol ^65^ and cholesterol is clustered at sites of protein secretion. ^66^ We hypothesised that the uptake of cholesterol driven by Scarf1 was being decoded by SNARE proteins to drive fibronectin secretion. The SNARE protein syntaxin 4 (Stx4) is the only cholesterol-sensitive SNARE protein present at the plasma membrane ^65^, and cholesterol promotes clustering of Stx4 with its partner SNAP23 at exocytic sites at the EC plasma membrane. ^65,67^ We silenced Stx4 expression (Figure SC) and examined the effect on LDL-induced fibronectin secretion. Silencing of Stx4 reduced LDL-stimulated apical fibronectin secretion (nLDL; 60% reduction, oxLDL; 47% reduction) when ECs were treated from the apical surface (Figure 7A). We repeated the experiment with lipoprotein treatment from the basal surface (Figure 7B). We found similar inhibition of basal fibronectin secretion (nLDL; 42% reduction, oxLDL; 40% reduction). We examined the cellular location of Stx4 in response to LDL treatment and saw translocation of Stx4 to the plasma membrane in ECs treated with either nLDL or oxLDL (Figure 7G).

**Figure 7.**
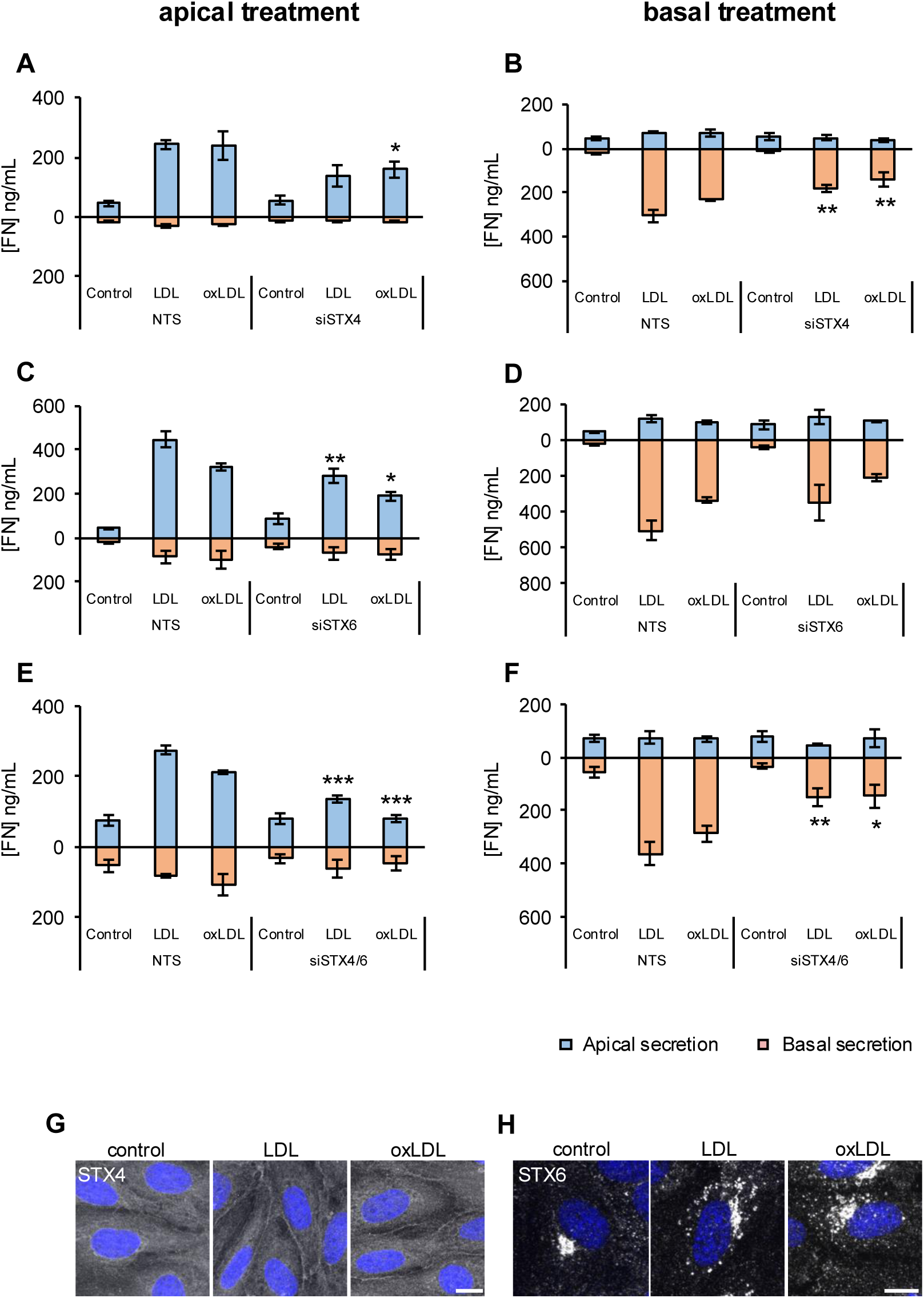
LDL-induced fibronectin secretion is regulated by syntaxin 4 and syntaxin 6. **A-F**, ECs were transfected with siRNAs targeting STX4 (**A,B**), STX6 (**C,D**) or both (**E,F**) and grown to confluence on Transwell inserts. Cells were treated with 50µg/mL nLDL or oxLDL from the apical or basal side as indicated. After 48h, the conditioned media were collected and the concentrations of apically-secreted (blue bars) or basally-secreted fibronectin (orange bars) were measured by ELISA and compared to the non-targeting siRNA controls (NTS). Data are means ± SEM; n = 3 experiments. **G,H,** ECs were grown to confluence on coverslips and treated with 50µg/mL nLDL or oxLDL for 48h. Cells were fixed and stained for **G,** STX4, and **H,** STX6. Plasma membrane translocation of Stx can be seen as a faint line around the border of the cells. Scale bars = 10 μm.

Silencing of Stx4 reduced lipoprotein induced fibronectin secretion by 40-60%, suggesting that other factors may also play a part. Syntaxin 6 (Stx6) is another cholesterol binding SNARE protein ^68^ and is involved in cholesterol trafficking from the late endosome to the trans-Golgi network (TGN). ^69^ Silencing of Stx6 expression effects endosomal recycling of the major fibronectin receptor integrin α5β1, ^65^ raising the possibility that it might be involved in fibronectin trafficking also. We silenced Stx6 expression (Figure S2D) and examined the effect on LDL-induced fibronectin secretion. Silencing of Stx6 reduced LDL-stimulated apical fibronectin secretion significantly (nLDL; 54% reduction, oxLDL; 65% reduction) when ECs were treated from the apical surface (Figure 7A). We repeated the experiment with lipoprotein treatment from the basal surface (Figure 7B). We found a lesser inhibition of basal fibronectin secretion (nLDL; 36% reduction, oxLDL; 46% reduction) and this did not reach statistical significance. We examined the cellular location of Stx6 in response to LDL treatment and saw translocation of Stx6 from the TGN into punctate structures resembling late endosomes in ECs treated with either nLDL or oxLDL (Figure 7H). We conclude that Stx6 is involved in cholesterol-mediated regulation of apical fibronectin secretion, acting at an internal trafficking step ahead of plasma membrane release.

Finally, we combined silencing of Stx4 and Stx6 to look for synergy of action. We now saw an almost complete block of apical fibronectin secretion in response to LDL treatment (Figure 7E; nLDL; 72% reduction, oxLDL; 99% reduction). We also saw increased inhibition of fibronectin secretion from the basal surface (Figure 7F). but this was still only partial inhibition (nLDL; 62% reduction, oxLDL; 51% reduction). Taken together, we conclude that the cholesterol-sensitive SNARE proteins Stx4 and Stx6 respond to cellular cholesterol changes caused by LDL uptake. Stx6 acts from an internal compartment to control onward trafficking of fibronectin to the surface, acting mostly in apical release. Stx4 mediates the secretory release of fibronectin at the EC surface, controlling release from both surfaces. In this way, binding of either nLDL or oxLDL to Scarf1 is coupled to re-routing of fibronectin intracellular trafficking and the consequent upregulation of fibronectin secretion from endothelial cells.

## DISCUSSION

The role of oxLDL as an inflammatory signal is well-studied. ^70^ In the early stages of atherosclerosis, nLDL accumulates in the subendothelial space. This becomes oxidised leading to activation of scavenger receptors, inflammatory signalling and the uptake of oxLDL by macrophages. ^47^ Our expectation was that any endothelial secretory response to LDL would be driven specifically by oxLDL and we were surprised to find that nLDL drove it equally. This suggests that this pathway functions as a response to levels of circulating cholesterol. In keeping with this, we found that the secretory response engages the cholesterol-sensitive SNARE proteins Stx4 and Stx6, allowing cholesterol to directly regulate the intracellular trafficking of the secreted proteins. Previous work has shown that oxLDL stimulates the release of extracellular vesicles (EVs) from endothelial cells. ^71^ Endothelial cells release EVs into the circulation to mediate long-range signalling. These membrane enclosed packages of chemokines, growth factors and micro-RNAS are then taken up by distant cells in the circulation to achieve a signalling outcome. ^9^ The secretory response that we have uncovered here would allow for more localised signalling in response to elevated LDL. The response to circulating LDL would result in release of pro-fibrotic proteins into the local circulation. Upon accumulation of LDL in the subendothelial space, the reversal of polarity would lead to release of those same proteins into the local vascular matrix.

We are used to thinking of LDLR as the receptor that mediates the uptake of cholesterol from nLDL, and scavenger receptors as providing inflammatory signals in response to oxLDL. We found that the endothelial secretory response was mediated by the less well-characterised scavenger receptor Scarf1, which was able to mediate uptake of cholesterol from both nLDL and oxLDL. Using a separate receptor in this way would allow the LDL-driven secretory response to function separately to the other pathways of normal cholesterol uptake, or the well-characterised responses to oxLDL.

The components of the secretory response cluster heavily on a set of processes, and the identities of these proteins give clues to the potential outcomes of this response. Many of the proteins have overlapping roles in fibrosis, atherosclerosis and senescence and they have multiple interconnections with each other. Further work needs to be done to examine how secretion of these proteins shapes disease progression; however, we can make some predictions at this point:

In the liver, sinusoidal ECs have a discontinuous basement membrane and contain fenestrae, allowing access of circulating LDL to the space of Diss where it is presented to the basal surface of the ECs. Our findings support a model whereby elevated circulating LDL would lead to basal secretion from the liver sinusoidal ECs, and the presentation of pro-fibrotic proteins to the adjacent hepatic stellate cells, promoting hepatic fibrosis.

And if we add this newly uncovered secretory response to what we already know of atherosclerosis, we get to a more complete picture of the endothelial response to LDL in that disease. Our findings support a model where elevated circulating nLDL leads to the apical release of proteins in the endothelial secretory response, sending pro-fibrotic, pro-inflammatory signals to the ECs in the local environment. If circulating LDL remains elevated over time, it starts to accumulate in the subendothelial space, presenting lipoprotein to the basal surface of the ECs. This would change the polarity of secretion and proteins like fibronectin would now be secreted into the vascular matrix, promoting ECM remodelling and endothelial to mesenchymal transition (Endo-MT). As this LDL became oxidised, the secretory response would be maintained, and we would proceed to previously characterised roles of oxLDL – engagement of scavenger receptors on the ECs to drive a transcriptional response, equipping the surface for leukocyte recruitment and amplifying oxidative and inflammatory stress. ^72^

In summary, this newly uncovered endothelial secretory pathway links circulating LDL to the release of pro-fibrotic proteins from the endothelium, supporting a role for endothelial cell secretion in lipid-induced fibrotic disease, and identifying Scarf1 as a potential target for pharmacological intervention.

## Non-standard Abbreviations and Acronyms

CCN1: Cellular Communication Network Factor 1
cFn: cellular fibronectin
DOC: deoxycholate
ECs: endothelial cells
Endo-MT: endothelial to mesenchymal transition ().
HUVEC: human umbilical endothelial cell
LDL: low-density lipoprotein, LDL receptor
nLDL: native LDL
oxLDL: oxidised LDL
LOX-1: lectin-like oxidized LDL receptor-1
MMP1: matrix metalloproteinase 1
PAI-1: Plasminogen activator inhibitor-1
Scarf1: Scavenger Receptor Class F Member 1
Stx: syntaxin
TMT: tandem mass tag.

## Acknowledgments

We would like to thank Martin Schwartz for early discussions on endothelial fibronectin secretion that set the course of the study. Exploratory work by Adelina Babii and Jonathan Furness supported the study but is not included. Stephen Cross from the Wolfson Bioimaging Facility provided support with fibronectin fibril quantification.

## Supplemental Material

Supplemental Methods

Tables S1–S2

Figures S1-S2

Major Resources Table

